# FERONIA and ANJEA do not have a conserved role in self-incompatible *Arabidopsis* for self-pollen rejection

**DOI:** 10.64898/2026.08.07.743519

**Authors:** Paula K.S. Chadic, Angela C. Sidsworth, Daphne R. Goring

## Abstract

The rejection of self-incompatible (SI) *Brassica* pollen is mediated by three signaling branches that function in parallel in the stigma. The recognition of SI pollen by the stigma S-Receptor Kinase (SRK) results in activation of the ARM-Repeat-Containing 1 E3 ubiquitin ligase (ARC1) which mediates the degradation of compatibility factors, the FERONIA (FER) and ANJEA (ANJ) receptor kinases that induces ROS accumulation to inhibitory levels and the *M* Locus Protein Kinase (MLPK) which may also be connected to ROS production. *Arabidopsis* self-incompatibility is regulated by SRK as well, but the signaling events downstream of SRK following SI pollen perception are less well-understood. In this study, we evaluated the requirements of *FER, ANJ* and *HERCULES RECEPTOR KINASE 1* (*HERK1*) for SI pollen rejection in the transgenic *Arabidopsis thaliana* SI-Col-0 *ψsrka-1* line. The *ψsrka-1* T-DNA disrupting the expression of the endogenous *ψSRKA* gene was crossed into SI-Col-0 to prevent any potential *SRK* transgene silencing. T-DNA mutants for *FER* and *ANJ/HERK1* were then crossed into the SI-Col-0 *ψsrka-1* line. Using standard assays for pollen-stigma interactions, the SI phenotypes were assessed for the SI-Col-0 *fer*, SI-Col-0 *anj-1* and SI-Col-0 *anj-1 herk1-1* lines. Our results presented here indicated that *FER* and *ANJ* are not required in the stigma for *Arabidopsis* SI pollen rejection, further providing evidence for a divergence in the SI downstream signaling pathway in *Arabidopsis*.

## Introduction, Results and Discussion

In self-incompatible (SI) Brassicaceae species, self-pollen is rapidly rejected following pollen contact with a stigmatic papilla at the top of the pistil through the actions of the tightly-linked *S* locus genes, *S Cysteine-Rich/S Protein 11* (*SCR/SP11*; hereafter *SCR*) and *S Receptor Kinase* (*SRK*). Both genes are polymorphic and specific *SCR-SRK* allele combinations define *S*-haplotype-specific interactions between the pollen SCR peptide and the stigma SRK leading to SI pollen rejection. The downstream signaling events following SCR-SRK mediated SI pollen rejection have been best described for *Brassica* and *Arabidopsis* species where the SRK signaling pathway in the stigmatic papilla blocks SI pollen hydration, an essential step for pollen germination (reviewed in (Goring et al., 2023; Nasrallah, 2023; Murase et al., 2024; Zhang et al., 2024). In *Brassica* SI, four signaling proteins act immediately downstream of activated SRK: the ARM-Repeat-Containing 1 E3 ubiquitin ligase (ARC1), the *M* Locus Protein Kinase (MLPK), and the FERONIA (FER) and ANJEA (ANJ) receptor kinases. There is no redundancy in these SRK-activated signaling components as the individual loss of ARC1, MLPK, FER or ANJ in the stigma causes a breakdown in SI pollen rejection (Stone et al., 1999; Murase et al., 2004; Chen et al., 2019; Zhang et al., 2021; Abhinandan et al., 2023; Huang et al., 2023). While some shared elements for SI signaling have been identified between *Brassica* and *Arabidopsis*, there are some unique features suggesting a divergence in the downstream signaling elements between these related species (reviewed in Goring, 2017; Jany et al., 2019; Goring et al., 2023). Our results presented here indicated that *FER* and *ANJ* are not required for *Arabidopsis* SI pollen rejection further providing evidence for this divergence in the SI downstream signaling.

*Arabidopsis thaliana* acquired mutations in *SCR, SRK* and *ARC1* during the transition to selfing, but the transformation of these genes from related *Arabidopsis* SI species into *A. thaliana* successfully restores the SI trait, and these transgenic lines have been used extensively to investigate the *Arabidopsis* SI signaling pathway (reviewed in Abhinandan et al., 2022; Goring et al., 2023; Nasrallah, 2023; Zhang et al., 2024). Interestingly, transforming *Brassica SCR* and *SRK* genes into *A. thaliana* failed to restore the SI trait, and the predicted MLPK ortholog, *Arabidopsis* APK1b, does not appear to be involved in *Arabidopsis* SI (Kitashiba et al., 2011; Zhang et al., 2019; Yamamoto et al., 2022). Also, there are accession differences for the requirement of *Arabidopsis* ARC1 to establish a strong SI phenotype (Boggs et al., 2009; Indriolo et al., 2012; Indriolo et al., 2014; Iwano et al., 2015).

Here we tested the requirement of FER and ANJ in *Arabidopsis* self-incompatibility using a transgenic *A. thaliana* SI line. This transgenic line carries the *Arabidopsis halleri SCR*^*13*^, *SRK*^*13*^ and *ARC1* genes on a single T-DNA construct in the Col-0 accession and displays a strong SI phenotype (Zhang et al., 2019; Macgregor et al., 2022; Zhang et al., 2024). As well, the *ψsrka-1* T-DNA disrupting the expression of the endogenous *ψSRKA* gene was crossed into this SI line to further stabilize the SI phenotype as *A. thaliana* Col-0 *ψSRKA* was shown to produce small RNAs that could silence related *SRK* transgenes (Fujii et al., 2020). The *fer-4* knockout and *fer-5* knockdown T-DNA mutants (Duan et al., 2010) and *anj-1* knockout T-DNA mutant (Galindo-Trigo et al., 2020) were then crossed into the transgenic *A. thaliana* Col-0 *SCR*^*13*^*-SRK*^*13*^*-ARC1 ψsrka-1* line (hereafter SI-Col-0) to evaluate their functions (all T-DNA identifiers and genotyping primers are listed in Supplementary Table S1). Since ANJ acts redundantly with HERCULES RECEPTOR KINASE 1 (HERK1) in a parallel function to *FER* in the synergid cells during pollen tube reception (Galindo-Trigo et al., 2020), HERK1 was also included in our analyses using the *herk1-1* knockout T-DNA mutant (Galindo-Trigo et al., 2020). The genotypes of all plants used in this study were confirmed by PCR genotyping (Supplementary Fig. S1 and Table S1). Furthermore, known phenotypes for the *fer* and *anj-1 herk1-1* mutants (Escobar-Restrepo et al., 2007; Duan et al., 2010; Galindo-Trigo et al., 2020) were confirmed in the SI-Col-0 homozygous mutant lines (Supplementary Fig. S2). Standard assays for pollen-stigma interactions were used to assess if *FER* and *ANJ/HERK1* were required in the stigma for SI pollen rejection (Supplementary Methods; Lee et al., 2020; Macgregor et al., 2022; Beronilla and Goring, 2024). First, pollen hydration assays were conducted using SI-Col-0 pollen to examine if the *fer, anj* or *anj herk1* mutations would affect the rejection of SI pollen at the critical pollen hydration step (Fig. 1, Supplementary Fig. S3). The control compatible pollination, SI-Col-0 pollen on Col-0 stigmas, displayed a large diameter increase resulting from hydration at 10-min post-pollination (Fig. 1a, c, d) while the SI-Col-0 pollen on the SI-Col-0 stigmas only showed a small diameter increase, typical of a SI pollen rejection phenotype at the hydration stage (Fig. 1b, c, d). The same phenotypes were observed for SI pollinations on the SI-Col-0 *fer-4* and SI-Col-0 *fer-5* stigmas (Fig. 1c), as well as the SI-Col-0 *anj-1* and SI-Col-0 *anj-1 herk1-1* stigmas (Fig. 1d). There were only small increases in SI-Col-0 pollen grain diameters and no significant difference to the control SI pollinations, with the exception of the SI-Col-0 *fer-5* (knockdown mutant) stigmas which supported a very small but significant increase over the control SI pollination. Thus, these results indicated that the mutations in *FER, ANJ* and *HERK1* fail to disrupt SI pollen responses in the SI-Col-0 stigmas at the hydration step. To confirm that this was a SI-specific phenotype and not a generalized hydration defect, compatible Col-0 pollen was used to pollinate the mutant SI-Col-0 lines (Fig 1c and d, Supplementary Fig. S3). As expected, Col-0 pollen was well-hydrated on all stigmas and displayed large increases in pollen diameter on the SI-Col-0 *fer-4* and SI-Col-0 *fer-5* stigmas (Fig. 1c) and the SI-Col-0 *anj-1* and SI-Col-0 *anj-1 herk1-1* stigmas (Fig. 1d). The average pollen grain diameters at 10-min post-pollination were not significantly different between control and mutant lines indicating that the *fer, anj* and *herk1* mutations did not impact the abilities of SI-Col-0 stigmas to hydrate compatible pollen. This confirmed that the block in pollen hydration on the SI-Col-0 stigmas is SI-specific and functions very effectively in the absence of *FER, ANJ* and *HERK1*.

**Fig 1.**
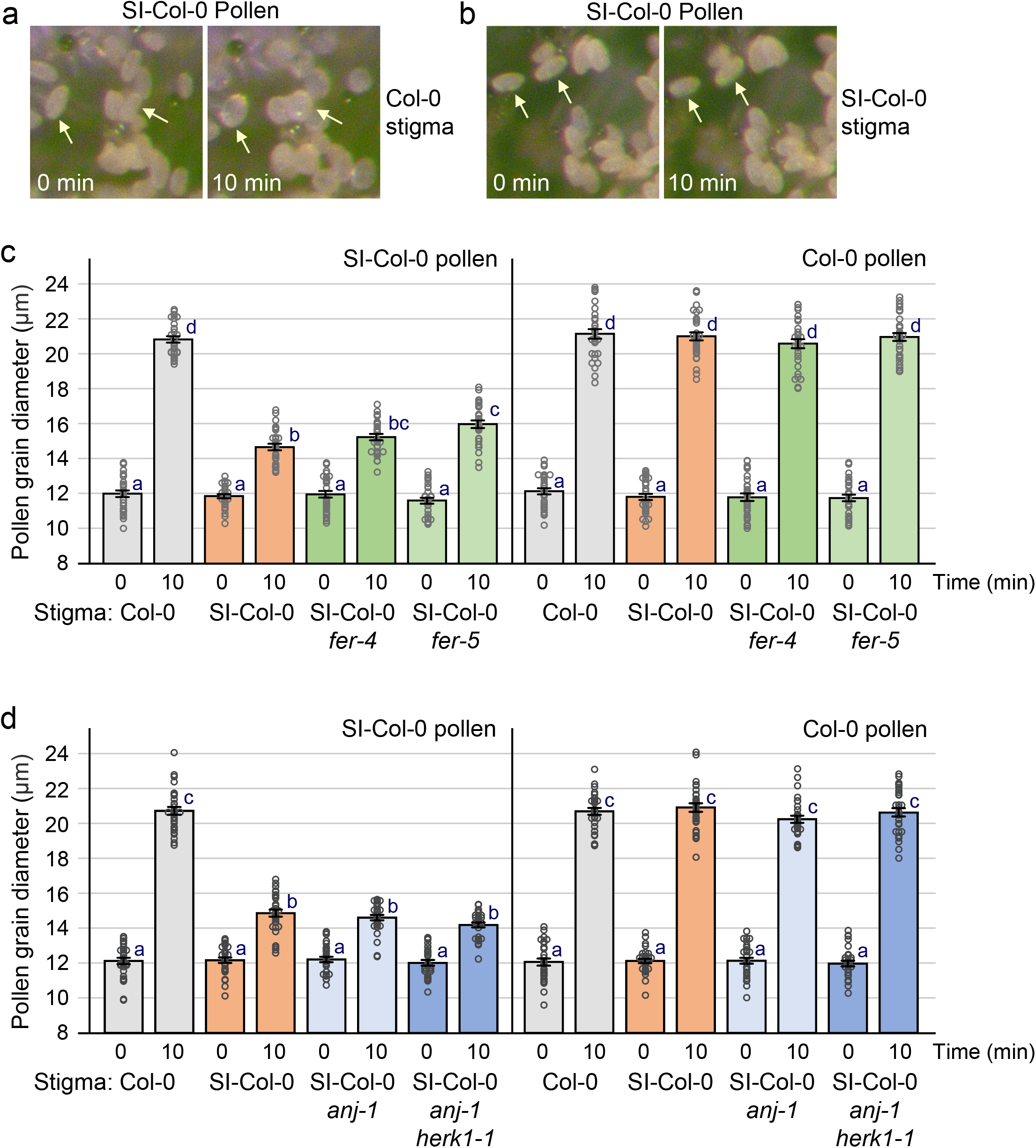
Pollen hydration assays show an intact SI pollen rejection response in SI-Col-0 mutant stigmas. **(a, b)** Images at 0 and 10-min post-pollination of a **(a)** control compatible pollen hydration and **(b)** control SI pollen hydration. Additional images are shown in Supplementary Fig. S3. **(c)**Pollen hydration assays on SI-Col-0 *fer-4* and SI-Col-0 *fer-5* mutant stigmas. Col-0 pollen and SI-Col-0 pollen were placed on stigmas from Col-0, SI-Col-0, SI-Col-0 *fer-4* and SI-Col-0 *fer-5* flowers. **(d)**Pollen hydration assays on SI-Col-0 *anj-1* and SI-Col-0 *anj-1 herk1-1* mutant stigmas. Col-0 pollen and SI-Col-0 pollen were placed on stigmas from Col-0, SI-Col-0, SI-Col-0 *anj-1* and SI-Col-0 *anj-1 herk1-1* flowers. As a proxy for hydration, pollen grain diameters were measured at 0- and 10-min post-pollination (Supplementary Methods; Lee et al. 2020). Data are shown as bar graphs of the means ± SE with all the data points displayed. *n* = 30 pollen grains per line. Letters represent statistically significant groupings of *P* <0.05 based on a one-way ANOVA with a Tukey-HSD post-hoc test.

Although most SI pollen grains are not hydrated sufficiently for germination, we did observe a few pollen grains forming pollen tubes in the SI-Col-0 line. Accordingly, we examine aniline blue stains for pollen tube growth in the mutant SI lines at 24h post-pollination. Overall, pollinations with SI-Col-0 pollen resulted in little to no pollen tubes in the SI-Col-0 *fer-4*, SI-Col-0 *fer-5*, the SI-Col-0 *anj-1* and SI-Col-0 *anj-1 herk1-1* stigmas (Fig. 2a, c, e; Supplementary Fig. S4). In contrast, Col-0 pollinations resulted in an abundance of pollen tube growth confirming that the *fer, anj* and *herk1* mutations did not impact compatible pollen tube growth (Fig. 2b, d, f; Supplementary Fig. S4). SI-Col-0 pollination of compatible Col-0 pistils also resulted in abundant pollen tube growth as expected (Fig. 2g, Supplementary Fig. S4). Furthermore, when the number of pollen tubes were counted, there were no significant differences between SI-Col-0 and the SI-Col-0 mutant lines in the average number of SI-Col-0 pollen tubes per style (Fig. 2h). These SI-Col-0 pollen tube counts were also significantly smaller than that observed for the compatible Col-0 pollinations of the SI-Col-0 and the SI-Col-0 mutant pistils (Fig. 2h). Altogether, the results presented by the aniline blue staining and the pollen tube counts support the SI Col-0 pollen hydration data that the *fer, anj* and *herk1* mutations do not alter the abilities of SI-Col-0 stigmas to reject SI pollen. All SI-Col-0 mutant lines displayed a strong SI phenotype and had no impact on SI pollen rejection in both stages observed. Thus, our results show that FER and ANJ/HERK1 are not required in the stigma for the *Arabidopsis* SI pathway, and this contrasts their defined roles in *Brassica* SI (Zhang et al., 2021; Huang et al., 2023).

**Fig. 2.**
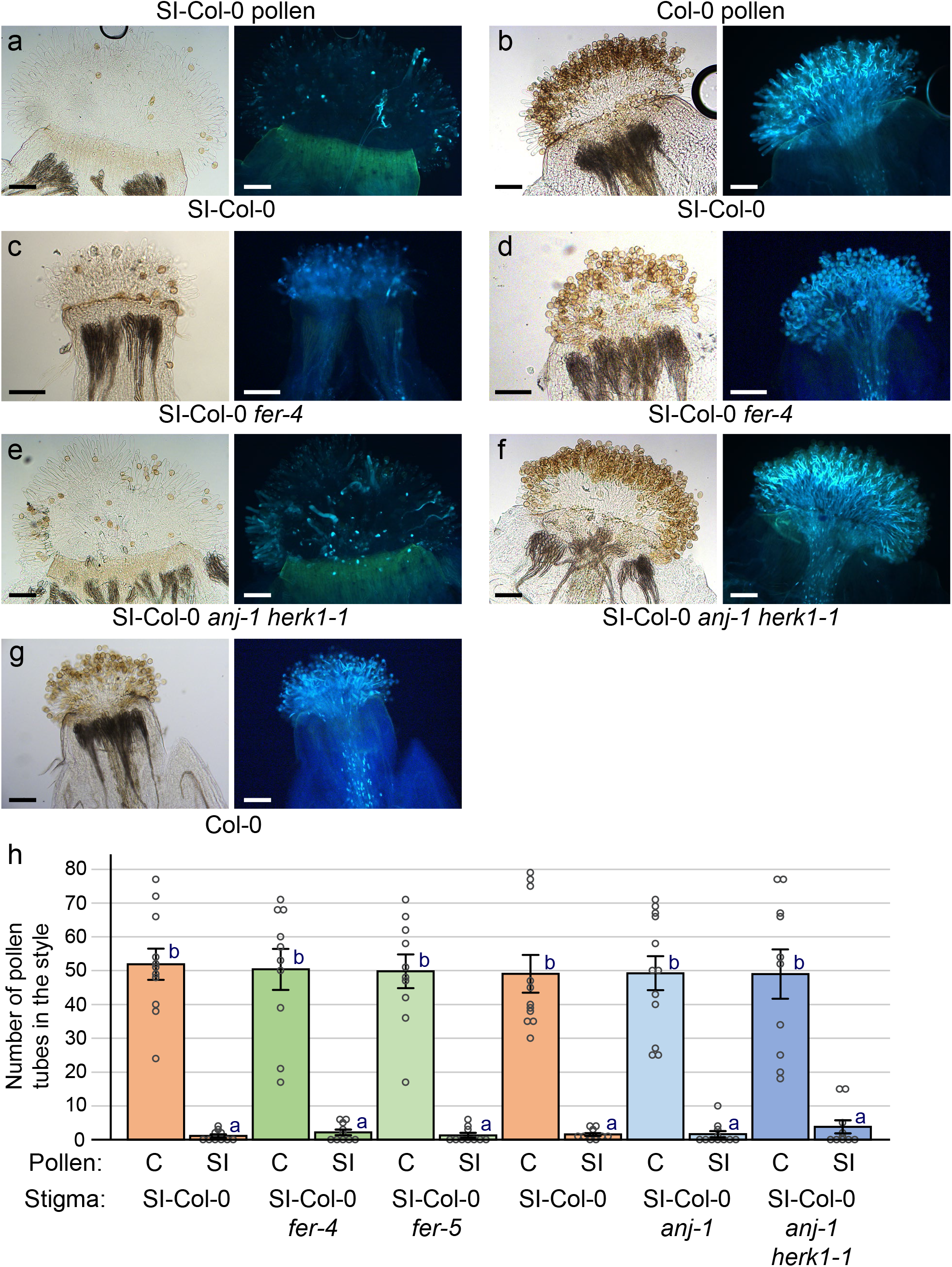
SI pollen rejection results in few pollen tubes in the SI-Col-0 mutant stigmas. **(a-g)** Representative images of aniline blue-stained SI-Col-0, SI-Col-0 *fer-4* and SI-Col-0 *anj herk1* stigmas collected at 24-h post-pollination. Pistils were pollinated with SI-Col-0 pollen or Col-0 pollen, and the genotypes of the pistils are indicated below. Brightfield (left) and aniline blue (right) images are shown for each sample. Pollen tubes growing through the pistil are indicative of pollen acceptance, as shown in panels **(b, d, f, g)** which represent compatible pollination controls. The fer, *anj* and *herk1* mutations did not have an impact on the stigma SI response as observed by the lack of SI-Col-0 pollen tube growth **(a, c, e)**. Scale bars = 100 µm. Additional images are shown in Supplementary Fig. S4. **(h)** Bar graph showing the mean number of compatible Col-0 (C) or SI-Col-0 (SI) pollen tubes/pistil at 24-h post-pollination. Data are shown as mean ± SE with all the data points displayed. *n* = 10-12 pistils per line. Letters represent statistically different groupings of *P* < 0.05 based on a one-way ANOVA with a Tukey-HSD post-hoc test.

With a focus on *Brassica* SI and unilateral incompatibility, Huang et al. (2023) reported key findings on the role of FER and ROS in these pathways, as well as confirming individual requirements of ARC1, MLPK, and ANJ in the stigma for SI pollen rejection. In addition, limited data was presented on an *Arabidopsis* SI-Col-0 *fer-4* mutant that was not consistent with the in-depth analyses presented here. While the same *Arabidopsis* SI-Col-0 line was used (Zhang et al., 2019), here the *ψsrka-1* T-DNA was also crossed into the SI-Col-0 line to prevent any potential SRK silencing (Fuiji et al 2020). Col-0 *ψSRKA* expresses a non-functional SRKA transcript with an exon 7 inverted repeat, producing sRNAs that can suppress related SRK transgenes, and the *ψsrka-1* T-DNA positioned in *ψSRKA* exon 7 disrupts sRNA production (Fujii et al., 2020). We observed that the presence of the *ψsrka-1* T-DNA in the SI-Col-0 had a more stabilizing effect on the SI phenotype with reduced age-related effects on the SI trait (Nasrallah et al., 2004), and this may account for the discrepancies between the two studies. An additional caveat is the potential epigenetic silencing of T-DNAs (Roca Paixao and Deleris, 2024), though silencing of the *SCR*^*13*^*-SRK*^*13*^*-ARC1* T-DNA would result in loss of the SI phenotype which we did not observe in this study.

Overall, our results show that FER and ANJ/HERK1 are not required in the stigma for the *Arabidopsis* SI pathway, providing additional evidence on the divergence of signaling events downstream of SRK signaling between *Brassica* and *Arabidopsis* (reviewed in Jany et al., 2019; Goring et al., 2023). While *Brassica* and *Arabidopsis* use the same SCR-SRK recognition system to initiate SI pollen rejection, transforming *Brassica SCR* and *SRK* genes into *A. thaliana* cannot restore the SI trait, except in some instances where a *Brassica* SRK kinase domain was replaced with an *Arabidopsis* SRK kinase domain (Zhang et al., 2019; Yamamoto et al., 2022).

Furthermore, the placement of a compatible pollen grain and an SI pollen grain on a single stigmatic papilla resulted in only the SI pollen rejected on the *Brassica* papilla, while both pollen grains were rejected on the *Arabidopsis* papilla, suggesting that *Brassica* SI has a more localized rejection response (Sarker et al., 1988; Dickinson, 1995; Iwano et al., 2015). Whether these observed differences are linked to the specific recruitment of FER in the *Brassica* SI pathway or novel downstream signaling proteins functioning in the *Arabidopsis* SI pathway will be key areas for future investigations.

## Acknowledgements

We thank work-study students (Cyrilla Zhang, Liz Lo, Flower Tan, Marina Kim, Gary Chatha, Hamna Ammar and Cecilia Widjaja) for their technical assistance, and Stuart Macgregor for assistance with the *A. thaliana* SI-Col-0 *ψsrka-1* crosses. We are also grateful to Alice Cheung (University of Massachusetts, Amherst) and Xue Pan (University of Toronto Scarborough) for the *fer-4* and *fer-5* mutant seeds, Lisa Smith (University of Sheffield) for the *anj-1 herk1-1* mutant seeds, and the ABRC for the *ψsrka-1* seeds (SALK_137645C).

## Author Contributions

PC and DG designed the research and wrote the first draft; PC and AS performed the research; All authors analyzed the data and edited the final version of the manuscript.

## Supplementary Data

The following materials are available in the online version of this article.

**Supplementary Methods**. Crosses, pollen hydration and aniline blue staining assays.

**Supplementary Figure S1**. Representative gel images for genotyping the SI-Col-0 mutant lines.

**Supplementary Figure S2**. Rosette photos and seed yields following compatible and SI pollinations.

**Supplementary Figure S3**. Representative Pollen hydration photos.

**Supplementary Figure S4**. Aniline blue stained images of SI-Col-0, SI-Col-0 *fer-4*, SI-Col-0 *fer-* SI-Col-0 *anj-1* and SI-Col-0 *anj-1 herk1-1* pistils pollinated with SI-Col-0 and Col-0 pollen.

**Supplementary Table S1**. Genotyping Primers.

## Data Availability

The data underlying this article are available in the article and in its online Supplementary data.

## Figure Legends

Fig 1. Pollen hydration assays show an intact SI pollen rejection response in SI-Col-0 mutant stigmas.

**(a, b)** Images at 0 and 10-min post-pollination of a **(a)** control compatible pollen hydration and**(b)** control SI pollen hydration. Additional images are shown in Supplementary Fig. S3.**(c)**

Pollen hydration assays on SI-Col-0 *fer-4* and SI-Col-0 *fer-5* mutant stigmas. Col-0 pollen and SI-Col-0 pollen were placed on stigmas from Col-0, SI-Col-0, SI-Col-0 *fer-4* and SI-Col-0 *fer-5* flowers.

**(d)** Pollen hydration assays on SI-Col-0 *anj-1* and SI-Col-0 *anj-1 herk1-1* mutant stigmas. Col-0 pollen and SI-Col-0 pollen were placed on stigmas from Col-0, SI-Col-0, SI-Col-0 *anj-1* and SI-Col-0 *anj-1 herk1-1* flowers. As a proxy for hydration, pollen grain diameters were measured at 0- and 10-min post-pollination (Supplementary Methods; Lee et al. 2020). Data are shown as bar graphs of the means ± SE with all the data points displayed. *n* = 30 pollen grains per line. Letters represent statistically significant groupings of *P* <0.05 based on a one-way ANOVA with a Tukey-HSD post-hoc test.

Fig. 2. SI pollen rejection results in few pollen tubes in the SI-Col-0 mutant stigmas. (a-g) Representative images of aniline blue-stained SI-Col-0, SI-Col-0 *fer-4* and SI-Col-0 *anj herk1* stigmas collected at 24-h post-pollination. Pistils were pollinated with SI-Col-0 pollen or Col-0 pollen, and the genotypes of the pistils are indicated below. Brightfield (left) and aniline blue (right) images are shown for each sample. Pollen tubes growing through the pistil are indicative of pollen acceptance, as shown in panels **(b, d, f, g)** which represent compatible pollination controls. The fer, *anj* and *herk1* mutations did not have an impact on the stigma SI response as observed by the lack of SI-Col-0 pollen tube growth **(a, c, e)**. Scale bars = 100 µm. Additional images are shown in Supplementary Fig. S4.

**(h)** Bar graph showing the mean number of compatible Col-0 (C) or SI-Col-0 (SI) pollen tubes/pistil at 24-h post-pollination. Data are shown as mean ± SE with all the data points displayed. *n* = 10-12 pistils per line. Letters represent statistically different groupings of *P* < 0.05 based on a one-way ANOVA with a Tukey-HSD post-hoc test.

## References

Abhinandan K, Hickerson NMN, Lan X, Samuel MA (2023) Disabling of ARC1 through CRISPR-Cas9 leads to a complete breakdown of self-incompatibility responses in Brassica napus. Plant Commun 4: 100504

Abhinandan K, Sankaranarayanan S, Macgregor S, Goring DR, Samuel MA (2022) Cell–cell signaling during the Brassicaceae self-incompatibility response. Trends in Plant Science 27: 472–487

Beronilla PKS, Goring DR (2024) Investigating a role for PUB17 and PUB16 in the self-incompatibility signaling pathway in transgenic Arabidopsis thaliana. Plant Direct 8: e622

Boggs NA, Nasrallah JB, Nasrallah ME (2009) Independent S-locus mutations caused self-fertility in Arabidopsis thaliana. PLoS Genet 5: e1000426

Chen F, Yang Y, Li B, Liu Z, Khan F, Zhang T, Zhou G, Tu J, Shen J, Yi B, Fu T, Dai C, Ma C (2019) Functional Analysis of M-Locus Protein Kinase Revealed a Novel Regulatory Mechanism of Self-Incompatibility in Brassica napus L. Int J Mol Sci 20

Dickinson H (1995) Dry stigmas, water and self-incompatibility in Brassica. Sexual Plant Reproduction 8: 1–10

Duan Q, Kita D, Li C, Cheung AY, Wu HM (2010) FERONIA receptor-like kinase regulates RHO GTPase signaling of root hair development. Proc Natl Acad Sci U S A 107: 17821–17826

Escobar-Restrepo JM, Huck N, Kessler S, Gagliardini V, Gheyselinck J, Yang WC, Grossniklaus U (2007) The FERONIA receptor-like kinase mediates male-female interactions during pollen tube reception. Science 317: 656–660

Fujii S, Shimosato-Asano H, Kakita M, Kitanishi T, Iwano M, Takayama S (2020) Parallel evolution of dominant pistil-side self-incompatibility suppressors in Arabidopsis. Nature Communications 11: 1404

Galindo-Trigo S, Blanco-Touriñán N, DeFalco TA, Wells ES, Gray JE, Zipfel C, Smith LM (2020) CrRLK1L receptor-like kinases HERK1 and ANJEA are female determinants of pollen tube reception. EMBO Rep 21: e48466

Goring DR (2017) Exocyst, exosomes, and autophagy in the regulation of Brassicaceae pollen-stigma interactions. Journal of Experimental Botany 69: 69–78

Goring DR, Bosch M, Franklin-Tong VE (2023) Contrasting self-recognition rejection systems for self-incompatibility in Brassica and Papaver. Current Biology 33: R530–R542

Huang J, Yang L, Yang L, Wu X, Cui X, Zhang L, Hui J, Zhao Y, Yang H, Liu S, Xu Q, Pang M, Guo X, Cao Y, Chen Y, Ren X, Lv J, Yu J, Ding J, Xu G, Wang N, Wei X, Lin Q, Yuan Y, Zhang X, Ma C, Dai C, Wang P, Wang Y, Cheng F, Zeng W, Palanivelu R, Wu H-M, Zhang X, Cheung AY, Duan Q (2023) Stigma receptors control intraspecies and interspecies barriers in Brassicaceae. Nature 614: 303–308

Indriolo E, Safavian D, Goring DR (2014) The ARC1 E3 ligase promotes two different self-pollen avoidance traits in Arabidopsis. The Plant Cell 26: 1525–1543

Indriolo E, Tharmapalan P, Wright SI, Goring DR (2012) The ARC1 E3 Ligase Gene Is Frequently Deleted in Self-Compatible Brassicaceae Species and Has a Conserved Role in Arabidopsis lyrata Self-Pollen Rejection. The Plant Cell 24: 4607–4620

Iwano M, Ito K, Fujii S, Kakita M, Asano-Shimosato H, Igarashi M, Kaothien-Nakayama P, Entani T, Kanatani A, Takehisa M, Tanaka M, Komatsu K, Shiba H, Nagai T, Miyawaki A, Isogai A, Takayama S (2015) Calcium signalling mediates self-incompatibility response in the Brassicaceae. Nat Plants 1: 15128

Jany E, Nelles H, Goring DR (2019) The Molecular and Cellular Regulation of Brassicaceae Self-Incompatibility and Self-Pollen Rejection. Int Rev Cell Mol Biol 343: 1–35

Kitashiba H, Liu P, Nishio T, Nasrallah JB, Nasrallah ME (2011) Functional test of Brassica self-incompatibility modifiers in Arabidopsis thaliana. Proc Natl Acad Sci U S A 108: 18173–18178

Lee HK, Macgregor S, Goring DR (2020) A Toolkit for Teasing Apart the Early Stages of Pollen–Stigma Interactions in Arabidopsis thaliana. In A Geitmann d, Pollen and Pollen Tube Biology: Methods and Protocols. Springer US, New York, NY, pp 13–28

Macgregor SR, Lee HK, Nelles H, Johnson DC, Zhang T, Ma C, Goring DR (2022) Autophagy is required for self-incompatible pollen rejection in two transgenic Arabidopsis thaliana accessions. Plant Physiol 188: 2073–2084

Murase K, Shiba H, Iwano M, Che F-S, Watanabe M, Isogai A, Takayama S (2004) A membrane-anchored protein kinase involved in Brassica self-incompatibility signaling. Science 303: 1516–1519

Murase K, Takayama S, Isogai A (2024) Molecular mechanisms of self-incompatibility in Brassicaceae and Solanaceae. Proc Jpn Acad Ser B Phys Biol Sci 100: 264–280

Nasrallah JB (2023) Stop and go signals at the stigma-pollen interface of the Brassicaceae. Plant Physiol 193: 927–948

Nasrallah M, Liu P, Sherman-Broyles S, Boggs N, Nasrallah J (2004) Natural variation in expression of self-incompatibility in Arabidopsis thaliana: implications for the evolution of selfing. Proceedings of the National Academy of Sciences 101: 16070–16074

Roca Paixao JF, Deleris A (2024) Epigenetic control of T-DNA during transgenesis and pathogenesis. Plant Physiol 197

Sarker RH, Elleman CJ, Dickinson HG (1988) Control of pollen hydration in Brassica requires continued protein synthesis, and glycosylation in necessary for intraspecific incompatibility. Proc Natl Acad Sci U S A 85: 4340–4344

Stone SL, Arnoldo M, Goring DR (1999) A breakdown of Brassica self-incompatibility in ARC1 antisense transgenic plants. Science 286: 1729–1731

Yamamoto M, Kitashiba H, Nishio T (2022) Generation of Arabidopsis thaliana transformants showing the self-recognition activity of Brassica rapa. Plant J 111: 496–507

Zhang D, Li Y-Y, Zhao X, Zhang C, Liu D-K, Lan S, Yin W, Liu Z-J (2024) Molecular insights into self-incompatibility systems: From evolution to breeding. Plant Communications 5: 100719

Zhang L, Huang J, Su S, Wei X, Yang L, Zhao H, Yu J, Wang J, Hui J, Hao S, Song S, Cao Y, Wang M, Zhang X, Zhao Y, Wang Z, Zeng W, Wu H-M, Yuan Y, Zhang X, Cheung AY, Duan Q (2021) FERONIA receptor kinase-regulated reactive oxygen species mediate self-incompatibility in Brassica rapa. Current Biology 31: 3004–3016.e3004

Zhang T, Wang K, Dou S, Gao E, Hussey PJ, Lin Z, Wang P (2024) Exo84c-regulated degradation is involved in the normal self-incompatible response in Brassicaceae. Cell Rep 43: 113913

Zhang T, Zhou G, Goring DR, Liang X, Macgregor S, Dai C, Wen J, Yi B, Shen J, Tu J (2019) Generation of transgenic self-incompatible Arabidopsis thaliana shows a genus-specific preference for self-incompatibility genes. Plants 8: 570

