## Supplementary Material for "FERONIA and ANJEA do not have a conserved role in self-incompatible *Arabidopsis* for self-pollen rejection"

### Supplementary Methods

#### Plant materials and growth conditions

All *A. thaliana* seeds were sterilized and stratified at 4$^{o}$C for 5 days in the dark. Stratified seeds were sown directly on Sunshine #1 soil supplemented with Plant Prod All Purpose 20-20-20 fertilizer for germination and growth. Plants were grown in growth chambers with growth cycles of 16-h light/8-h dark at 22$^{o}$ C. Humidity in the chambers were maintained below 50%.

#### Crossing SI-Col-0 lines with the mutant lines

The transgenic *A. thaliana* SI-Col-0 lines #2 and #3 (carrying *AhSCR-AhSRK-AhARC1* transgenes, Zhang et al., 2019) were crossed with the *ψsrka-1* mutant (Fujii et al. 2020) to generate the *A. thaliana* Col-0 $SCR_{13}$-$SRK_{13}$-*ARC1 ψsrka-1* line (abbreviated to SI-Col-0). These plants were then crossed with the *fer-4*, *fer-5*, and *anj-1 herk1-1* mutants (Duan et al., 2010; Galindo-Trigo et al., 2020) to produce the different *A. thaliana* SI-Col-0 mutant lines. Two different *fer* mutants were tested: *fer-4* which is a knockout mutant and *fer-5* which is a knockdown mutant (Duan et al., 2010). The *fer-4* and *fer-5* mutations were successfully crossed into the *A. thaliana* SI-Col-0 line #3, while the *herk1-1* and *anj-1* mutations were successfully crossed into the *A. thaliana* SI-Col-0 line #2. Each plant used in this study was rigorously genotyped before analyzed in the pollen-pistil analyses. All T-DNA identifiers and genotyping primers are listed in Supplementary Table S1, and representative genotyping gels are shown in Supplementary Figure S1.

#### Pollen hydration and aniline blue staining assays

All post-pollination assays were performed as previously described (Lee et al., 2020; Macgregor et al., 2022; Beronilla and Goring, 2024). As outlined in Lee et al. (2020), the relative humidity was monitored with hygrometers in the growth chambers and lab so that experiments (i.e. pollen hydration assays) were performed when the relative humidity was less that 40% in the growth chambers and lab. All assays began with the emasculation of wildtype and mutant stage 12 flower buds. Emasculated pistils were wrapped in plastic wrap, and the plants were stored in the growth chambers. For the pollen hydration assay, emasculated pistils were extracted from the plants at 24-h post-emasculation and mounted onto ½ MS plates. A single anther from an open flower was utilized to lightly pollinate the mounted pistils. Images were captured through the Nikon sMz800 microscope at 0- and 10-min post-pollination. 10 pollen grains/stigma were randomly selected and measured using the NIS-elements imaging software. A total of 30 pollen grains were measured per group (3 pistils per line; $n$ =10/pistil). For the aniline blue staining assay, emasculated pistils were lightly pollinated with a single anther from an open flower, and pollinated pistils were collected 24-h post-pollination for aniline blue staining as described in Lee et al. (2020). Aniline blue stained pistils were mounted on slides with sterile water, flattened with a coverslip and imaged at 10X magnification on a Zeiss Axioscope2 Plus fluorescent microscope. Images were captured using the brightfield and UV fluorescence settings on the fluorescent microscope.

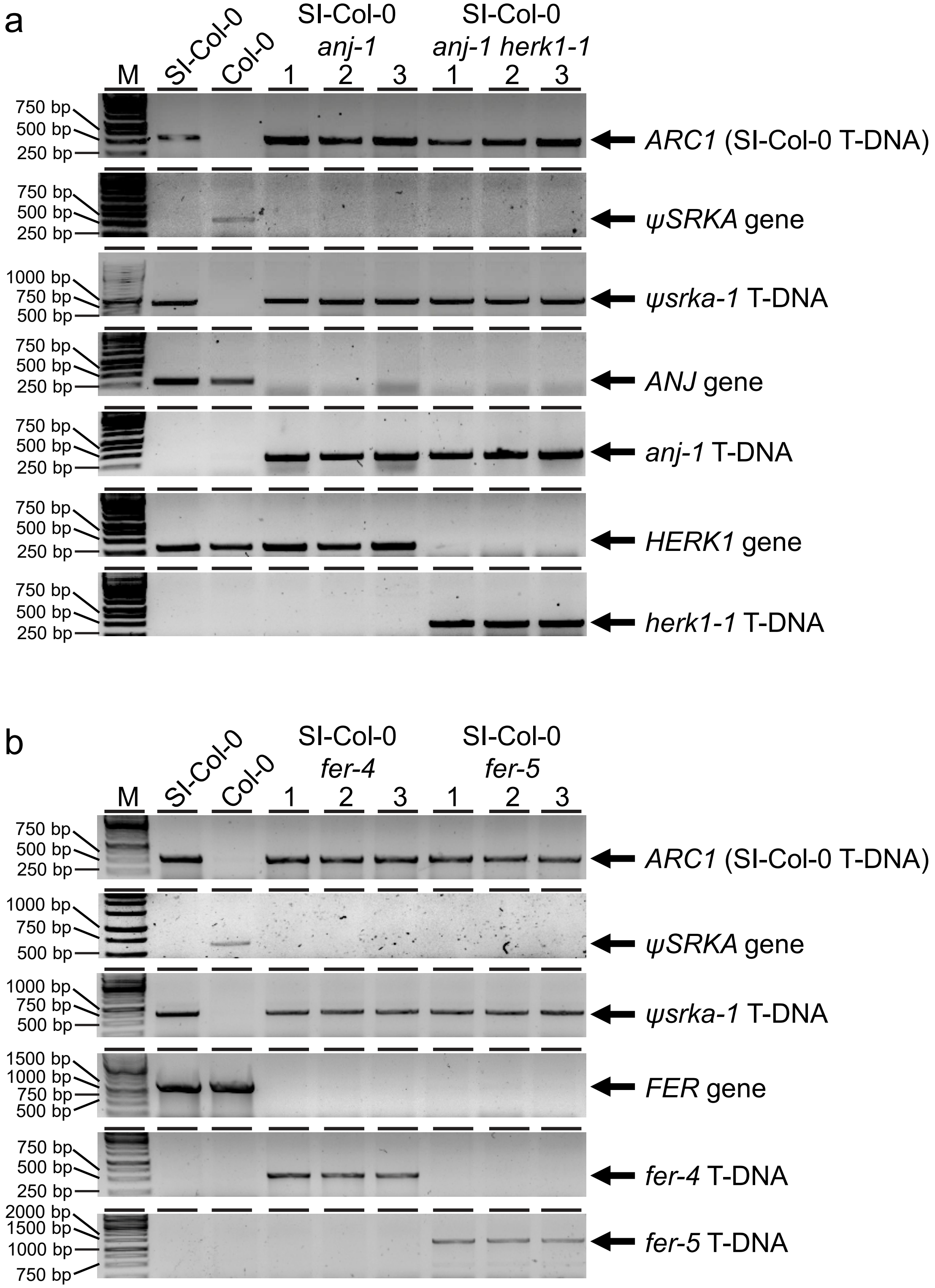


**Supplementary Figure S1. Representative gel images for genotyping the SI-Col-0 mutant lines.**
**(a)** Genotyping gels for the SI-Col-0, Col-0, SI-Col-0 *fer-4*, and SI-Col-0 *fer-5* lines as labeled at the top.
**(b)** Genotyping gels for the SI-Col-0, Col-0, SI-Col-0 *anj-1*, and SI-Col-0 *anj-1 herk1-1* lines as labeled at the top.
Primers used for each gene or T-DNA mutant are listed in Supplementary Table S1.

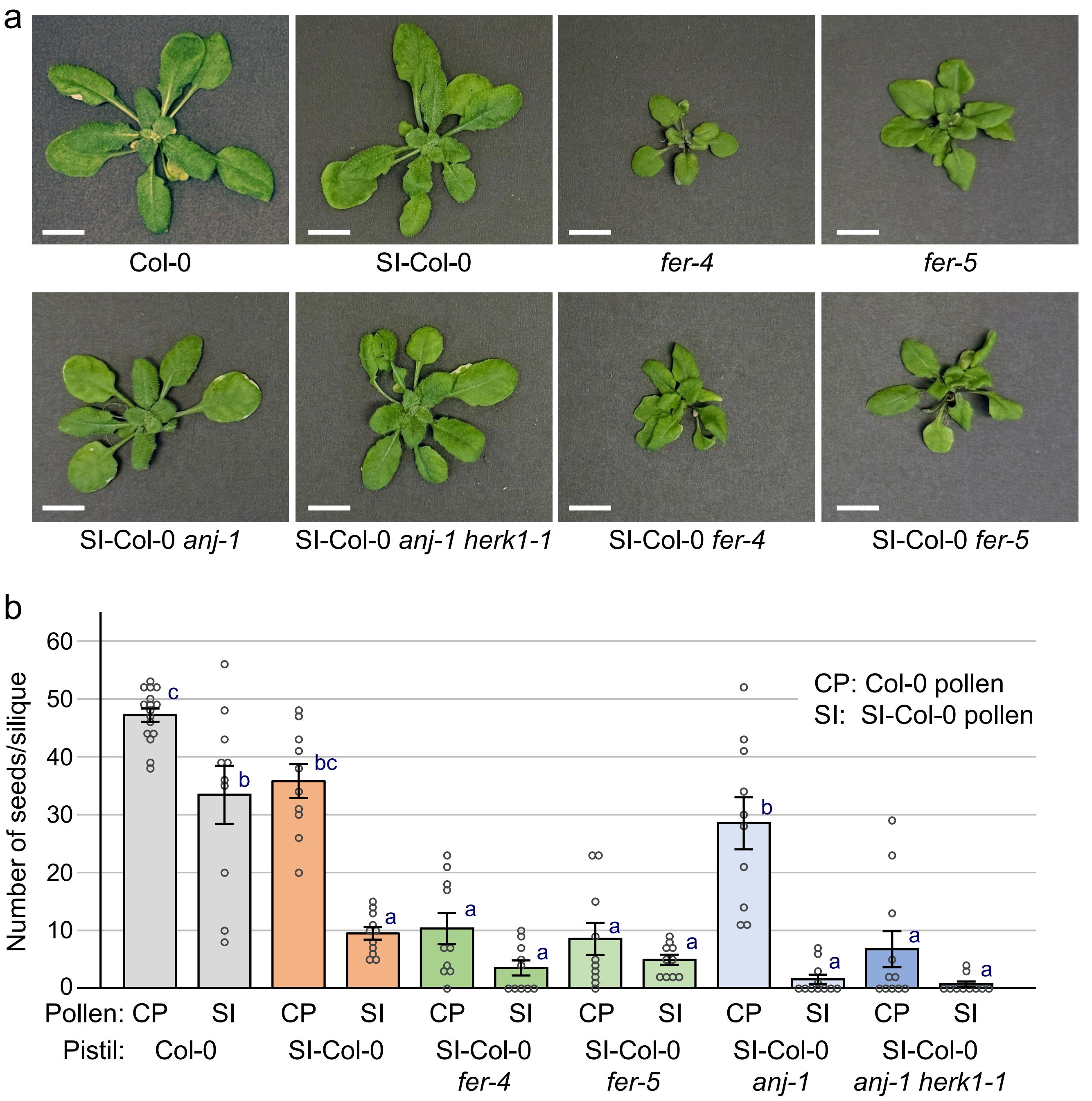


**Supplementary Figure S2. Rosette photos and seed yields following compatible and SI pollinations.**

**(a)** Representative photos of 4-week-old rosettes. The *fer* and SI-Col-0 *fer* mutant plants display the smaller rosette sizes expected for the *fer* mutants. See Duan et al. (2010) and Galindo-Trigo et al. (2020) for comparative images. Scale bars = 1 cm.

**(b)** Seed yields following pollinations of pistils from Col-0, SI-Col-0, SI-Col-0 *fer-4*, SI-Col-0 *fer-5*, SI-Col-0 *anj-1* and SI-Col-0 *anj-1 herk1-1* flowers with Col-0 (CP) or SI-Col-0 (SI) pollen. Siliques were collected at 2-weeks post-pollination for clearing and counting. The SI-Col-0 *fer* and the SI-Col-0 *anj-1 herk1-1* siliques display the expected reduced seed yield with compatible Col-0 due to the downstream pollen tube reception defect (Escobar-Restrepo et al., 2007; Galindo-Trigo et al., 2020). See Galindo-Trigo et al. (2020) for comparative data.

Data are shown as a bar graph of the means ± SE with all data points displayed. $n$ = 10-11 siliques per line. Letters represent statistically significant groupings of $P < 0.05$ based on a one-way ANOVA with a Tukey-HSD post-hoc test.

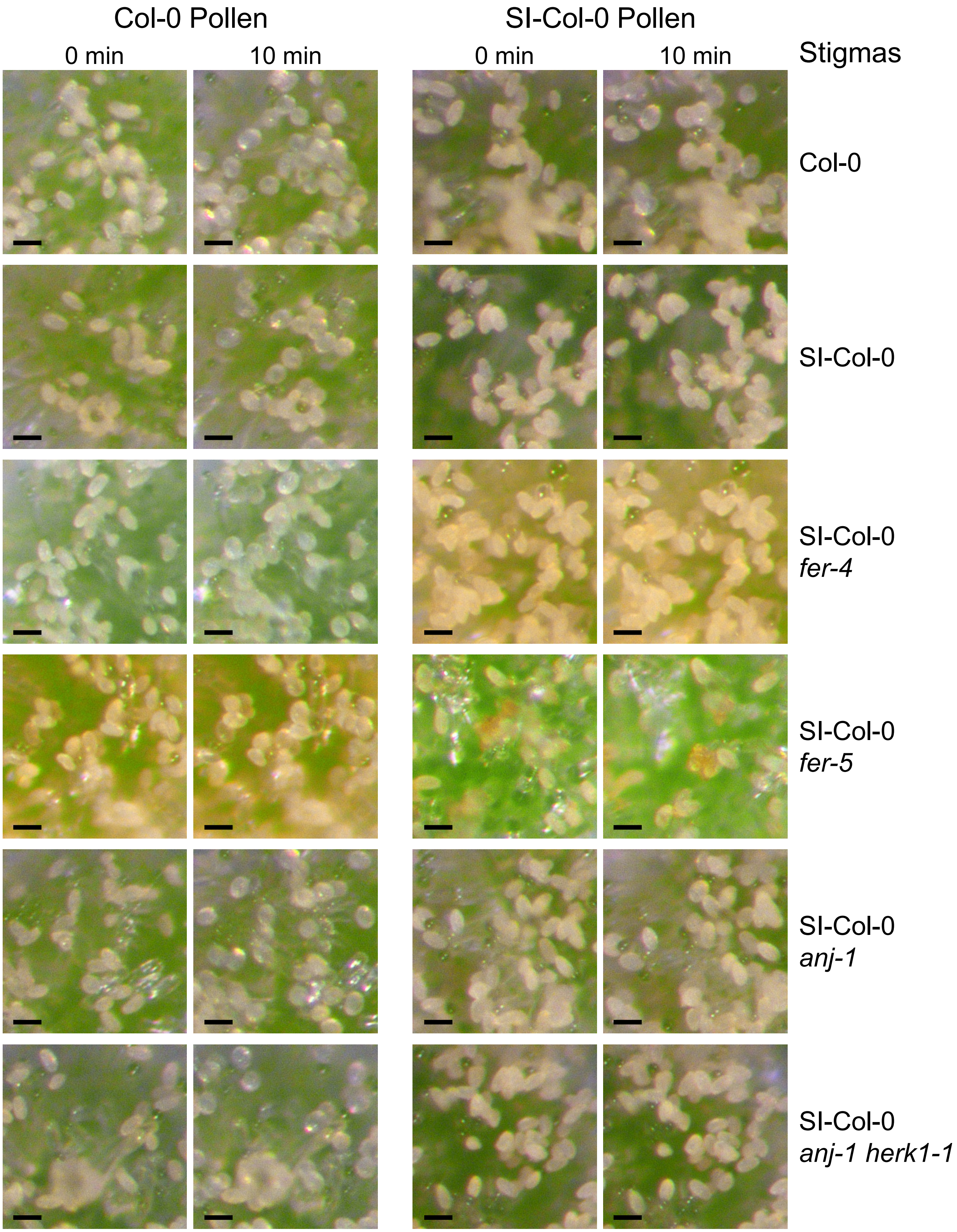


**Supplementary Figure S3. Representative Pollen hydration photos.**
See Fig.1 for the accompanying bar graphs. Scale bars = 25 μm.

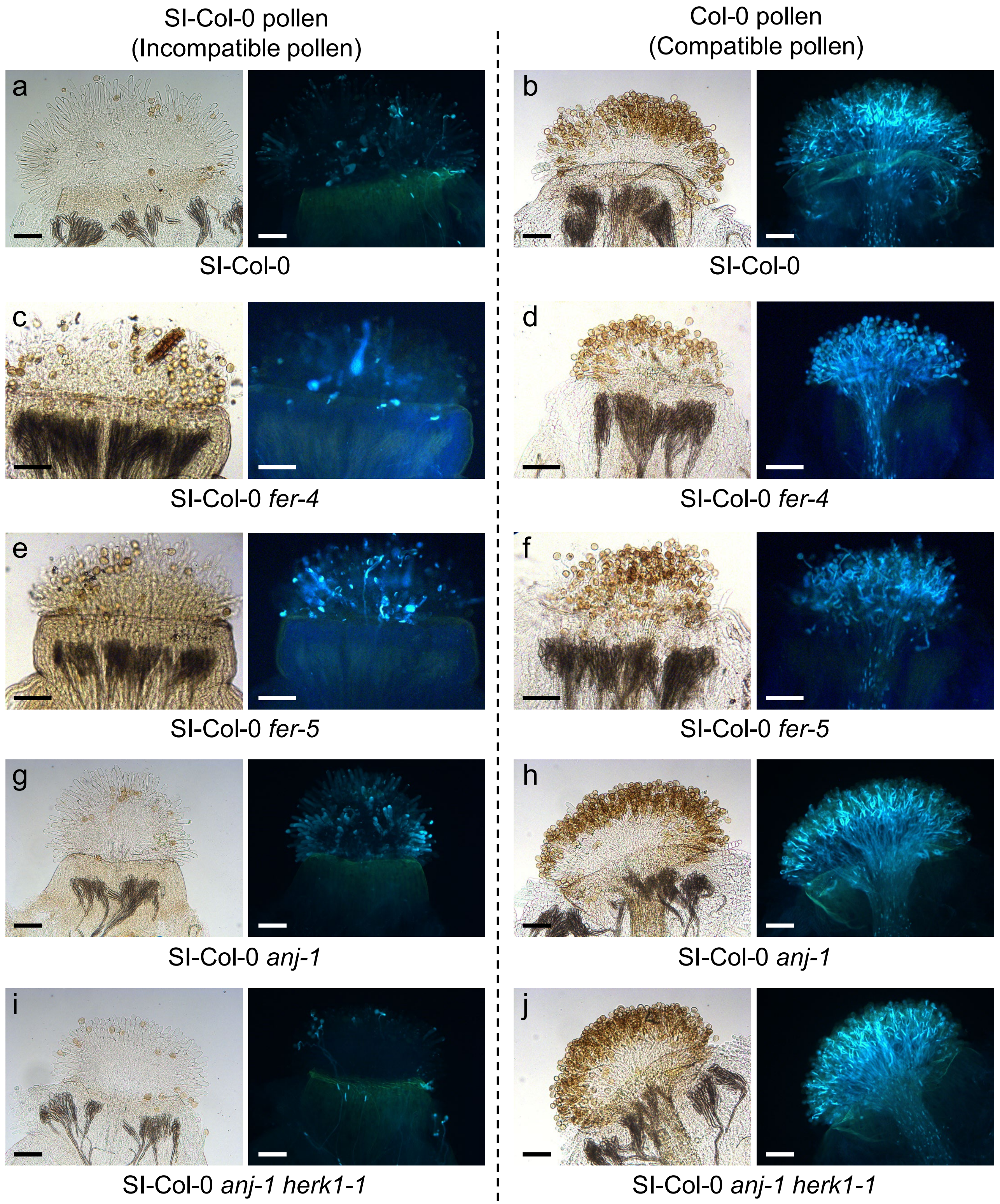


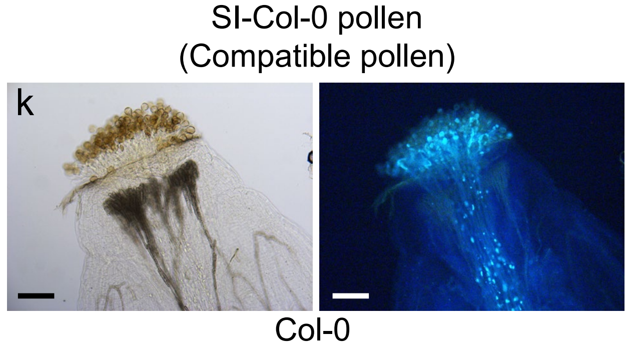


**Supplementary Figure S4. Aniline blue stained images of SI-Col-0, SI-Col-0 *fer-4,* SI-Col-0 *fer-5,* SI-Col-0 *anj-1* and SI-Col-0 *anj-1 herk1-1* pistils pollinated with SI-Col-0 and Col-0 pollen.** Representative images of pistils collected at 24-h post-pollination for aniline blue staining. Pistils were pollinated with incompatible SI-Col-0 pollen **(a, c, e, g, I)** or compatible Col-0 pollen **(b, d, f, h, j)**. **(k)** is a control to show that SI-Col-0 pollen is compatible on the Col-0 stigma. The genotypes of the pistils are indicated below, and brightfield (left) and aniline blue (right) images are shown for each sample. Scale bars = 100 µm.

### Supplementary Table S1. Genotyping Primers

| Genotyping Function | | Primer sequences |
|---|---|---|
| AhARC1 in the AhSCR-SRK-ARC1 T-DNA | FP | CCGAAGTGCTACAGAGTGGA |
| (in SI-Col-0) Zhang et al., 2019 | RP | TCCGAATCAGAGAACGTGAG |
| wildtype ψSRKA allele (At4g21370) | RP | GCTAACACCAGGAAGGTGGTTGG |
| Fujii et al., 2020 | FP | CCGTTTTGTACTCGTATGACG |
| ψsrka-1 T-DNA insertion (SALK_137645C) in | LBb1.3 (T-DNA) | ATTTTGCCGATTTCGGAAC |
| SRKIR$^{Col-0}$ (Col-0 ψSRKA) Fujii et al., 2020 | RP | TGACAATTGCATTCAAAAGGG |
| wildtype ANJEA allele (At5g59700; | RP | CTCCTCTGTAGCAAAACCAGGA |
| Galindo-Trigo et al., 2020 | FP | CTCACGTTTACTCCCTCGGG |
| anj-1 T-DNA insertion (SALK_114667C) | LBb1.3 (T-DNA) | ATTTTGCCGATTTCGGAAC |
| Galindo-Trigo et al., 2020 | FP | CTCCTCTGTAGCAAAACCAGGA |
| wildtype HERK1 allele (At3g46290) | FP | GTTGCTCGCGGTAGTCTTCT |
| Galindo-Trigo et al., 2020 | RP | CTGTCCTGAATTCCGCAAGC |
| herk1-1 T-DNA insertion (SALK_008043C) | LBb1.3 (T-DNA) | ATTTTGCCGATTTCGGAAC |
| Galindo-Trigo et al., 2020 | RP | CTGTCCTGAATTCCGCAAGC |
| wildtype FERONIA allele (At3g51550) | FP | CGGATCCATGGCTTACCGCAGACGTAAGCGTGG |
| Duan et al., 2010 | RP | CGAATTCACGTCCCTTTGGATTCATCATCTG |
| fer-4 T-DNA insertion (GABI_GK106A06) | LP | CGGATCCATGAAGATCACAGAGGGACGATTC |
| Duan et al., 2010 | GAB1-R (T-DNA) | GTGGATTGATGTGATATCTCC |
| fer-5 T-DNA insertion (SALK_029056) | LP | CGGATCCATGGCTTACCGCAGACGTAAGCGTGG |
| Duan et al., 2010 | LB1 (T-DNA) | GCGTGGACCGCTTGCTGCAACT |
